# Infraslow dynamic patterns in human cortical networks track a spectrum of external to internal attention

**DOI:** 10.1101/2024.04.22.590625

**Authors:** Harrison Watters, Aleah Davis, Abia Fazili, Lauren Daley, TJ LaGrow, Eric H. Schumacher, Shella Keilholz

**Author notes:** **Corresponding authors:** Shella Keilholz. Eric H. Schumacher. **Author Contributions:** Conceptualization, data preparation, data processing, formal analysis, figure drafting, paper writing, review, editing: HW, AD, AF. Data visualization, analysis, review, and editing: LD. Data preparation and curation: ES. Supervision, review, and editing: SK, ES. Preprocessing troubleshooting, review, and editing: TJL. **Competing Interest Statement:** The authors declare they have no competing interests.

## Abstract

Early efforts to understand the human cerebral cortex focused on localization of function, assigning functional roles to specific brain regions. More recent evidence depicts the cortex as a dynamic system, organized into flexible networks with patterns of spatiotemporal activity corresponding to attentional demands. In functional MRI (fMRI), dynamic analysis of such spatiotemporal patterns is highly promising for providing non-invasive biomarkers of neurodegenerative diseases and neural disorders. However, there is no established neurotypical spectrum to interpret the burgeoning literature of dynamic functional connectivity from fMRI across attentional states. In the present study, we apply dynamic analysis of network-scale spatiotemporal patterns in a range of fMRI datasets across numerous tasks including a left-right moving dot task, visual working memory tasks, congruence tasks, multiple resting state datasets, mindfulness meditators, and subjects watching TV. We find that cortical networks show shifts in dynamic functional connectivity across a spectrum that tracks the level of external to internal attention demanded by these tasks. Dynamics of networks often grouped into a single task positive network show divergent responses along this axis of attention, consistent with evidence that definitions of a single task positive network are misleading. Additionally, somatosensory and visual networks exhibit strong phase shifting along this spectrum of attention. Results were robust on a group and individual level, further establishing network dynamics as a potential individual biomarker. To our knowledge, this represents the first study of its kind to generate a spectrum of dynamic network relationships across such an axis of attention.

**Significance Statement:** This study provides significant insight into how cortical dynamics shift along a spectrum of external vs internal attention. Association and sensory networks show shifts in their relationship with the default mode network (DMN) along this spectrum. Notably, the constituent networks of the task positive network (TPN), such as the DAN, frontoparietal network (FPN), and ventral attention network (VAN) show divergent responses along this spectrum, indicating that definitions of a single “task positive” network are misleading. This is one of the first studies of large-scale spatiotemporal dynamics across such a breadth of tasks. These results provide a neurotypical reference spanning an axis of attention to interpret the growing functional dynamics literature and to help characterize functional dynamics as a potential biomarker.

## Introduction

The human cerebral cortex, which, to quote Geschwind & Rakic, “considers itself the crowning achievement of evolution”, is marked by significant expansion compared to other mammalian brains (1), with the majority of volume increase accounted for by disproportionate representation of non-sensory association cortex (2-4). Expansion of the association cortex paralleled the evolution of highly interconnected brain networks that became untethered to direct sensory stimuli and adapted novel functions related to attentional demand, memory, cognition, and social interaction (2).

These flexible but distinct cortical networks (5), originally detected in positron emission tomography (PET) and later functional magnetic resonance imaging (fMRI) (6, 7), largely maintain their organization at rest, in the absence of any externally directed task (8, 9). One way to view these distributed cortical networks is along an axis of external to internal attention (2, 10-12). At one end of the spectrum, regions of the default mode network (DMN) become less active during externally focused tasks, and more active during memory recall, mind-wandering, the maintenance of internal thought trains, and other internally directed processes (6, 7, 13-20). At the other end of the spectrum, regions showing task related activation, opposite the DMN, have been referred to as the task positive network (TPN), which typically is considered to include the dorsal attention network (DAN) (21, 22), the frontoparietal network (FPN)(11), and sometimes the ventral attention or salience network (VAN) (23, 24). The DMN seems to work in opposition with networks that exhibit increased activity during externally directed tasks, showing anticorrelated activity with the DAN specifically (10, 25-28). As measured with fMRI when subjects are at rest, anticorrelated activity between the DMN and DAN appears to be a hallmark of neurotypical cognition. DMN-DAN anticorrelation declines in aging and dementias (29-34). Diminished DMN-DAN anticorrelation has also been seen in sleep deprivation (35, 36), is associated with attention deficit hyperactivity disorder (ADHD) in children (37, 38), poor task performance in adults (39), and the use of exogenous neuromodulators that negatively affect working memory performance, such as delta9-tetrahydrocannabinol (40-42).

Anticorrelated DMN-DAN activity may have evolved as an adaptation to allocate cortical resources based on a spectrum of external vs. internal attention, and to allow for more elaborate construction of internal thoughts and narratives (10, 43-46). Weber et al reports that subjects engaged in a congruence task showed DMN deactivation that scaled with the difficulty of the task (47), indicating a relationship between task difficulty and increasing allocation of cognitive resources. In other words, when the external task was easier, the default mode network could afford to become more active. This interpretation of DMN-DAN anticorrelation fits with anecdotal human experience: when unrelated thoughts interrupt external tasks, we tend to perform the tasks poorly. Conversely, when external stimuli are overwhelming or distracting, maintaining an internal train of thought is difficult. Importantly, this model of network resource allocation also implies that underlying networks such as the DMN may show dynamic changes related to external vs internal attentional demand.

Seminal fMRI studies of cortical network activity, including DMN-DAN anticorrelation, relied on traditional time-averaged measures of activity, which represent the averaged correlation between reference-informed regions across the entire scan. While this time-averaged analysis approach is incredibly valuable for exploring external vs internal attention, it misses recurring dynamic patterns that capture time-varying relationships between networks within the fMRI signal (48, 49). A subject’s attention may shift repeatedly over the course of a single scan, and time-averaged approaches fail to capture network dynamics that may correspond to such attentional fluctuations. Across imaging modalities, there is increasing evidence that dynamic spatiotemporal patterns are related to arousal, attention, and cognition (50-60). Several types of dynamic spatiotemporal patterns have been described in fMRI literature (57, 61-64). The work in the current study focuses on the quasi-periodic pattern (QPP): a repeating, whole brain spatiotemporal pattern of alternating activity between the DMN, TPN, and other networks that happens quasi-periodically on the infraslow timescale (lasting about 20 seconds in humans), and has been directly implicated in attention and arousal (50, 53, 55, 60, 65, 66). Consistent with previously mentioned time-averaged FC studies regarding the DMN, QPPs are disrupted in individuals with ADHD (54) and low frequency waves of DMN activity (<0.1Hz) disrupt task performance (67). More recently, work from collaborators showed that the amount of anticorrelation in the QPP between DMN and the frontoparietal network (FPN) decreases along with task performance (59).

Considering the important evolutionary role of the DMN, its critical functions in attention and cognition, and the potential for inter-network DMN dynamics as a biomarker of healthy brain function across task and rest states, we tested whether the dynamic relationships of networks in infraslow QPPs reflect the axis of external vs. internal attention. Here, we detected QPPs in a range of datasets (Figure 1) obtained from a mix of open sources and collaborators (Supplemental table 1). We employed inter-network correlation analysis (Figure 3) between the DMN and DAN, as well as all other major attentional networks of the cortex (Figure 2) during the QPP. We hypothesized that 1) key attentional networks would show task related shifts in their correlation with the DMN. 2) These effects would be robust when QPPs were detected at both the group averaged and individual level. 3) The constituent networks of the so-called task-positive network (i.e., DAN, FPN, VAN) would show divergent results with respect to their correlation with the DMN in the QPP. And finally, 4) sensory cortices would also participate in dynamic shifting of DMN correlation in response to the task-rest spectrum.

**Table 1.** Datasets and QPP parameters.

| Dataset | OpenNeuro<br>VWM | VG:<br>Gamer<br>Task | HCP<br>Task/<br>Rest | Flanker | Simon | Stroop | CABI<br>Rest | Meditators | Twilight<br>Zone |
| --- | --- | --- | --- | --- | --- | --- | --- | --- | --- |
| EPI TR (s) | 2.53 | 0.535 | .72 | 2.5 | 2 | 1.5 | 2.25 | 1.5 | 1.5 |
| QPP<br>Window<br>Length | 8 | 32 | 30 | 8 | 10 | 12 | 10 | 10 | 12 |
| Final QPP<br>Length (s) | 20.24 | 17.12 | 21.6 | 20 | 20 | 18 | 22.5 | 15 | 18 |

**Figure 1.**
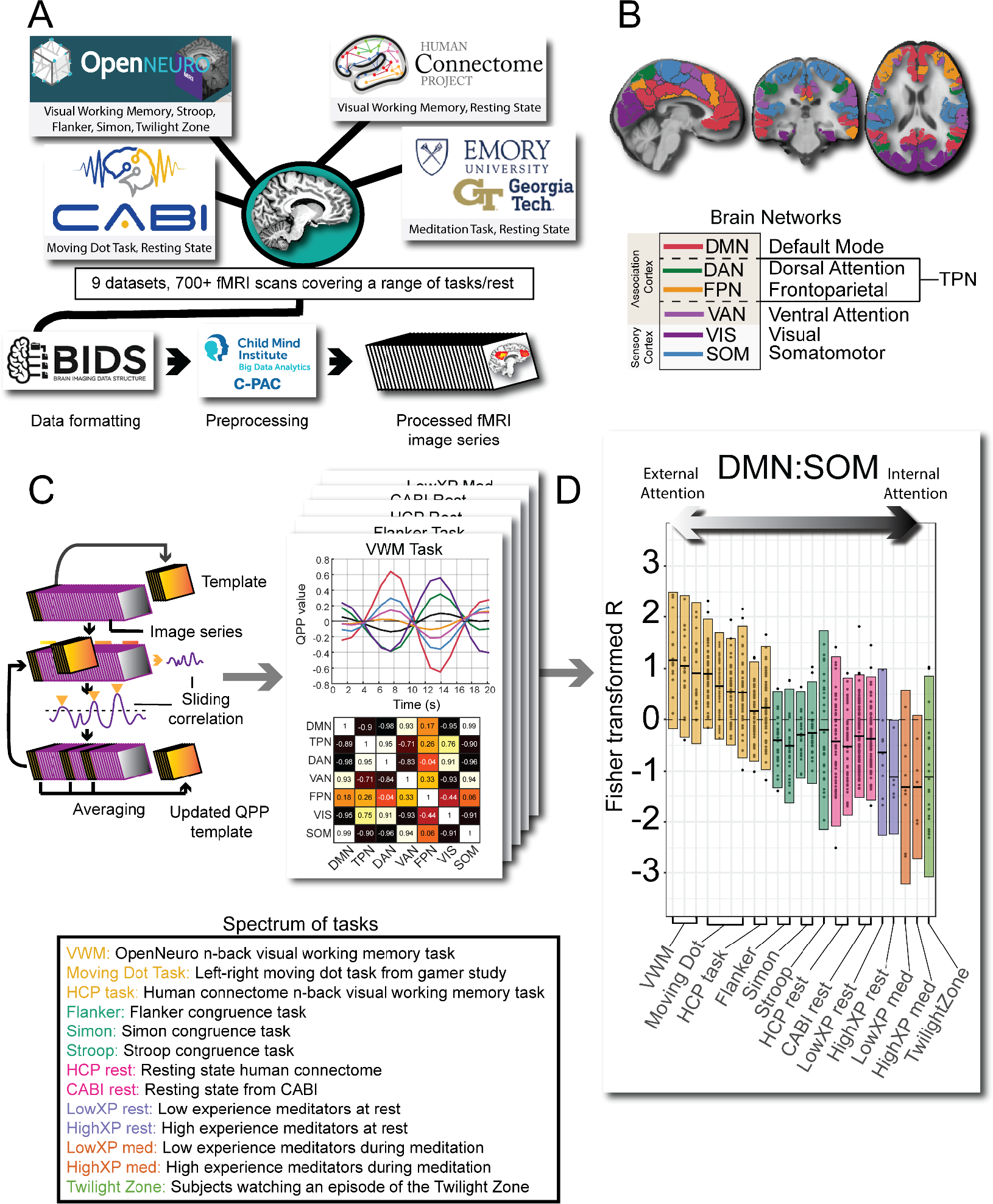
(A) Data sources to CPAC processing. (B) Example of Yeo’s network used in the present study, shown in a 3-plane slice of standard MNI 152 space. Note that the task positive network (TPN) is the combination of DAN and FPN networks. (C) Process of QPP detection, network waveforms, and inter-network correlations. (D) Example of inter-network analysis specific to DMN. Note the change in correlation to the DMN across an axis of external to internal attention.

**Figure 2.**
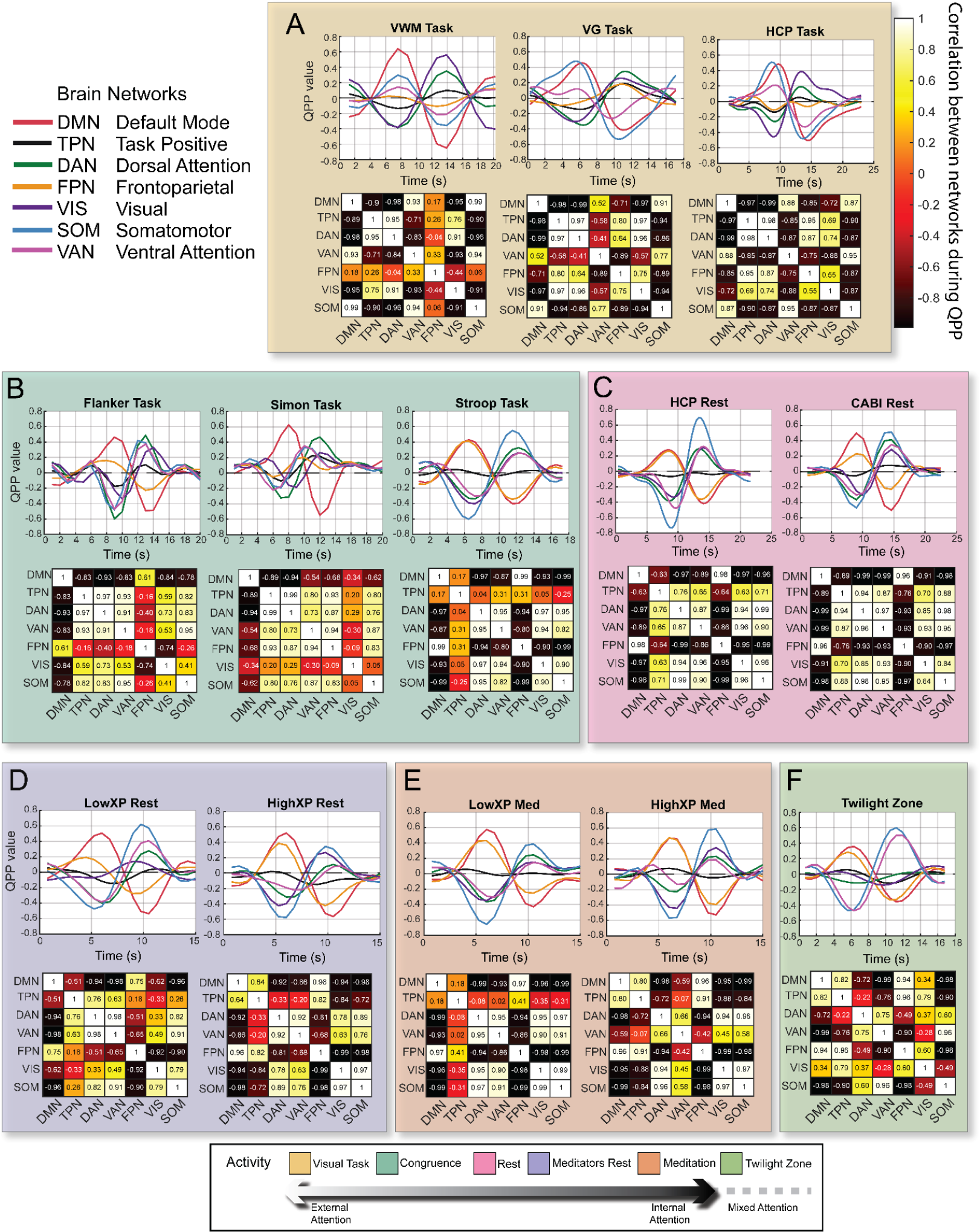
(A-F) QPP waveforms and heatmaps showing correlations between all networks during QPP across task spectrum. Note switching of SOM and VAN from negative DMN correlation at rest/congruence to positive DMN correlation in visual tasks. FPN becomes more negatively correlated with DMN in visual and congruence tasks compared to rest.

**Figure 3.**
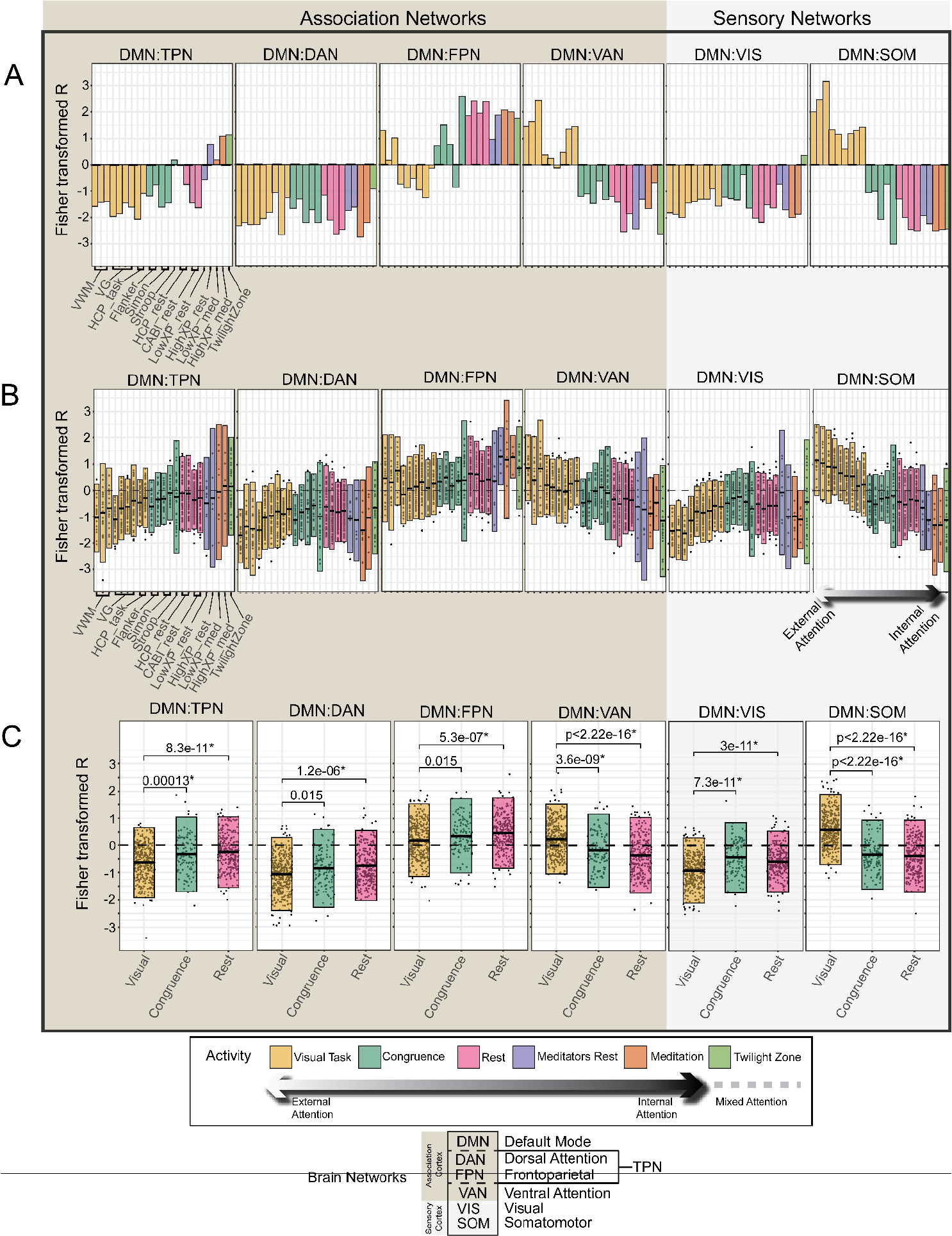
Shifts in DMN-to-network correlations along the spectrum of fMRI tasks, shown as Fisher transformed correlation R values on y-axis vs task type on x-axis. Group (A), Individual (B), and Stat Comparisons (C). Note that DAN, FPN, and VIS networks show task-positive-like responses moving from rest towards visual tasks (right to left), while VAN and SOM show the opposite trend, becoming positively correlated with the DMN during visual tasks.

This is one of the first studies connecting such a breadth of datasets for spatiotemporal analysis in relation to an axis of attention. The results provide significant theoretical insight into the role of inter-network DMN dynamics in the balance of external vs internal attention. Additionally, the resulting dynamic DMN fingerprints for each task type can be used as a reference to guide future studies, to interpret disparate findings related to dynamic functional connectivity, and to help identify shifts in inter-network DMN activity to be used as a biomarker in health and disease.

## Results

Our results reflect the dynamic relationships of cortical networks within QPPs, spatiotemporal patterns of activity, expressed as the correlation between networks (for example movies of such QPPs see other work by our group (55, 56). The reported correlation structure simply summarizes the behavior of specific networks within these patterns. Overall, QPPs showed a strong pattern of DMN/DAN anticorrelation, consistent with previous literature, using both seed-based and dynamic FC methods, describing strong anticorrelation between default and task positive brain networks (10, 17, 18, 53, 55, 56, 59, 60). Group level QPP waveforms and inter-network group correlations are reported first followed by subject-wise DMN QPP correlations and statistical comparisons. For both group and individual analysis, we find that a spectrum of inter-network DMN shifting emerges based on task type that seems to track a spectrum of external to internal attention.

### 3.1 Group level QPP waveforms and qualitative network differences across task type

Several major qualitative differences and trends were quickly apparent in the group level QPP waveforms (Figure 3). First, in the visually demanding tasks (moving dot task and visual working memory, see methods), the phase of the VAN and somatosensory (SOM) networks is completely switched compared to the other task types (compare Figure 3A to other task types). That is, rather than showing positive correlation with the DAN they are instead strongly correlated with the DMN. This VAN and SOM phase switching is not apparent in congruence tasks, though they do trend more positively in DMN correlation (Figures 2 and 3A).

The FPN also shows a major shift corresponding to the spectrum of tasks (Figure 2 and Figure 3A-B). Specifically, across all resting groups and during meditation and TV watching the FPN is strongly in phase with the DMN. For both visual and congruence tasks the FPN becomes more positively correlated with the DAN than the DMN. The more externally directed the current task is (further left on task spectrum in Figure 3A), the stronger the FPN-DAN alignment in the QPP appears to be on average. This is consistent with previous seed-based FC findings characterizing the frontoparietal control network as a flexible network that can shift between modulating the DAN or DMN, respectively, depending on task type (43, 68).

The DMN-TPN relationship at the group level is somewhat more variable across the datasets, trending towards positive in the meditation and TV watching datasets, and more negative correlation in the visual task sets. The increased DMN-TPN anticorrelation during tasks found here is consistent with literature indicating that externally focused tasks increase separation between the DMN and TPN (58). However, as mentioned, the TPN described in this study and in others is just the combination of the DAN with other attentional networks such as the FPN and VAN, which show divergent results in our QPP data with respect to their DMN correlation.

When inspecting the results of the DAN and FPN across the task and rest spectrum reported in the present study, it is apparent that shifts in activity of the FPN are primarily responsible for the overall TPN inter-network correlation shift. In the group level QPP, the DMN-DAN correlation is universally negative with minimal variability, only trending towards the positive in the Twilight Zone set. The FPN, however, shows dynamic switching in the QPP that corresponds strongly to the amount of externally directed task demand the subjects are engaged in (Figure 3A). In other words, in the group level QPP, the DAN exhibits only small change in DMN anticorrelation based on task, where the FPN seems highly responsive to the task-rest spectrum, shifting flexibly between the DMN at rest and the DAN during externally directed tasks.

### 3.2 Individual level QPP correlations for Visual, Congruence, and Rest states

Subject-wise inter-network DMN correlation results were largely consistent with group level findings (**Figure 3 A-B**). Specifically, means of the individual DMN-to-network correlations for all networks showed the same trends found at the group level. That is, in both group and individual QPPs the TPN, DAN, FPN, and VIS showed significantly decreased DMN correlation during visual tasks compared to rest and congruence, while the VAN and SOM showed significantly increased DMN correlation for visual tasks compared to rest and congruence (**Figure 3C**). In summary, both group and individual results indicate that with respect to infraslow QPP dynamics, the VAN and SOM exhibit task-negative-like responses, shifting phase to be more in line with the DMN, a network associated with internal mentation, memory retrieval, mind-wandering, and perceptual decoupling (6, 16, 18, 28, 45). Meanwhile, the DAN, FPN, and VIS networks exhibit task-positive-like responses.

It is important to highlight the results of the HCP dataset within the task-rest spectrum, as they act as a reference point to validate the other datasets analyzed here. Most of the datasets have subjects with either a task or a rest scan, not both. For example, the CABI rest subjects were only scanned during rest, and the VG and VWM subjects were only scanned during tasks. The HCP subjects however were each scanned twice at rest and twice during a visual working memory task (69). Thus, the HCP subjects plotted in the task and rest categories are the same subjects. Importantly, these same subjects compared at task and rest recapitulate the same results of network phase shifting and correlation changes for each DMN-to-network comparison made between tasks in the other datasets (note HCP task and rest results in Figure 3A-B).

### 3.3 Meditators and Twilight Zone DMN correlations during the QPP

While meditators and Twilight Zone groups contained much smaller sample sizes and thus were not included in the statistical comparison with the visual, congruence, and rest groups, they exhibited notable trends that inform the other datasets. The high experience meditation group, actively meditating subjects, and the Twilight Zone group trended much higher than resting state subjects in their DMN-DAN correlation, at the opposite extreme from external visual tasks. A similar but opposite trend is seen in the DMN-SOM correlation, with the high experience meditation group, actively meditating subjects, and the Twilight Zone group showing a much more negative DMN correlation, resembling extreme rest. For networks such as the DAN, VAN, and SOM, these miscellaneous datasets (meditators/Twilight Zone) suggest that activities related to a high degree of internal focus or narration cause inter-network DMN shifts that fit on the resting state end of the attention spectrum. Or rather, that tasks involving heavy internal focus or narration overlap heavily in their inter-network DMN dynamics with resting states.

### 3.4 Inter-network DMN fingerprints during QPP

Inter-network DMN correlations for all networks were co-plotted on polar plots to capture a functional connectivity fingerprint for each task state (Figure 4). The overall fingerprints are notably similar between congruence tasks and rest (Figure 4), while the visual task varies significantly (captured in Figures 2/4 qualitatively and statistically in network-by-DMN comparisons in Figure 3C). Meditation and Twilight Zone DMN fingerprints also trend strongly away from typical rest. In conjunction, the results of inter-network DMN dynamics during the QPP in Figures 3 and 4 indicate a spectrum of network shifting with respect to the DMN that corresponds heavily to the balance of external vs internal attention in each task.

**Figure 4.**
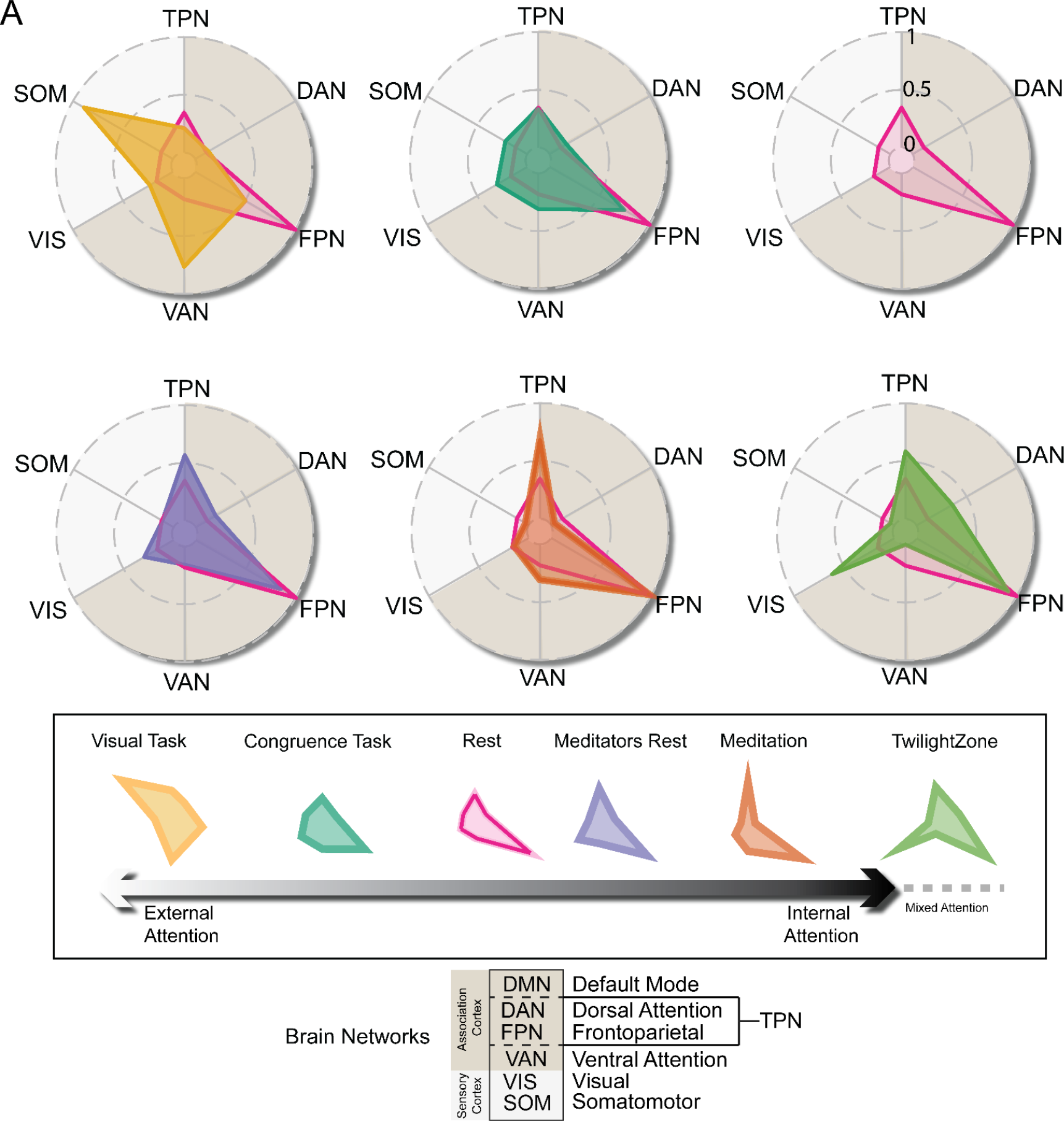
DMN-to-network dynamics for each category of task, plotted as correlation to DMN for each network normalized to a scale of 0 (minimum) to 1 (maximum). Note that all tasks are co-plotted with rest as a reference state. Correlational structure for each task may represent a dynamic fingerprint for whole brain dynamics corresponding to the mix of external vs internal attention demanded by a task.

## Discussion

Here, we generated a spectrum of inter-network DMN dynamics from infraslow QPPs across a range of task types in neurotypical human subjects. Most whole brain spatiotemporal analysis projects advanced by our group and others have focused on single datasets and only on the relationship between the DMN and TPN. To our knowledge, this represents the first study of its kind to generate a spectrum of dynamic network relationships covering such a breadth of tasks and networks. These results show the value of the whole brain analysis compared to seed based or static FC measures. The approach of whole brain spatiotemporal pattern detection resolved time-varying changes of the locus of attention within network dynamics which would not have been detected with traditional time-averaged measures. We propose that the detected shifts in spatiotemporal network patterns tracks an axis of external to internal attention, with certain networks exhibiting dynamic task-positive-like behavior (DAN, FPN, VIS) and others showing task-negative-like dynamic shifting (DMN, VAN, SOM) during the QPP. This work has broad implications for the interpretation of dynamic FC findings and their implementation in preclinical research. For example, these neurotypical network dynamic results can be used as a reference as we and other groups apply QPP analysis to datasets of pathology and neurodivergence, helping us to identify biomarkers and understand how aberrant network dynamics contribute to the etiology of neural disorders. Additionally, these findings contribute important evidence to theory regarding the evolution of flexible cortical networks and attentional resource management. The robustness of the inter-network DMN shifting across group and individual level QPP analysis also indicates that QPPs may be a viable biomarker for individual subjects, not only group averaged level analysis. We discuss below interpretations of these findings for major networks as well as the potential use of these QPP dynamic fingerprints in guiding related studies and as a biomarker in health and disease.

### 4.1 Changes to attentional/association networks (DAN/FPN/VAN) across the task-rest spectrum

#### Dorsal attention network

We found significantly increased DMN-DAN anticorrelation during visual tasks. Notably, our QPP results indicate that definitions of a single “task positive” network can be misleading, given the divergent task responses shown by the DAN, FPN, and VAN, depending on the type of task and degree of external attention required (c.f., (55, 59). Our results are consistent with traditional seed-based FC evidence. Increased functional connectivity within the DAN is associated with improved performance and learning in visual tasks (70), and increased DMN-DAN anti-correlation is likely a feature of healthy cognition and a predictor of task performance (38). Additionally, our QPP results are corroborated by recent work from collaborators. Employing QPP analysis to a finger tapping task, Seeburger and colleagues found consistent results with respect to DMN-DAN anticorrelation. Subjects engaged in the finger tapping task were assessed for time spent “in the zone” vs “out of the zone”. Subjects who performed better showed significantly greater DMN-TPN anticorrelation, with some of this effect accounted for by increased DMN-DAN anticorrelation (59). Also consistent with our results, Seeburger found that the shifting in TPN anticorrelation was primarily driven by changes in the amount of FPN anticorrelation, not DAN, once again indicating that use of the overarching “task positive” network definition can obscure important divergence in underlying network behavior.

#### Frontoparietal control network (also referred to as executive control network)

We found that the FPN exhibits decreased DMN correlation in the infraslow QPP depending on the task, becoming more positively correlated with the DMN during rest and more anticorrelated with the DMN during congruence and visually demanding tasks. This result lines up with a considerable body of evidence from fMRI and seed-based analysis that the FPN, or frontoparietal control network, acts to flexibly modulate the DMN and DAN depending on the degree of internal to external attention involved in a task (43, 68, 71-76). For example, Smallwood and colleagues proposed that increased DMN-FPN alignment during rest, when attention is often directed inward or to mind-wandering, is critical for buffering internal thought trains to prevent interruption by intruding thoughts or external stimuli (43-45, 77, 78). Morin and colleagues, which employed the same Yeo’s 7 network definitions used in the present study, also reported flexible reconfiguration of the FPN resulting in increased DAN-FPN alignment and decreased DMN-FPN alignment during symbolic visual tasks (76). The FPN has also been further parcellated into subregions based on preferential connectivity with either the DMN and DAN, with these FPN subdivisions showing flexible change in DMN FC based on task type (74).

While FC studies of meditation offer mixed results (79-81), experienced practitioners of mindfulness meditation, an activity marked by repeatedly practicing executive control of one’s attention, have also shown increased FC between the DMN-FPN, an effect reported using seed-based and sliding window measures of FC (80, 82). In meditators included in the current study (83), we found DMN-FPN correlation trended more positive both at rest and during meditation, an effect seen in group and individual QPPs (Figure 3A-B). In seeming contradiction with the previous studies, Godwin et al reported that subjects with a high propensity for mind-wandering showed increased resting state FC between the DMN and FPN (84).This seems at odds with DMN-FPN alignment driving more executive control of thought trains and buffering from mind-wandering. However, the intentionality of mind-wandering may explain this disparity, as intention may have a major impact on DMN-FPN interaction (85, 86). Golchert et al found that subjects engaged in intentional mind-wandering showed increased DMN-FPN FC compared to those experiencing unintentional mind-wandering (85). There is also some evidence that decreased DMN-FPN FC at rest is correlated with higher IQ scores and part of normal brain maturation (87).

While there are disparate results and many open questions regarding the role of DMN-FPN connectivity in neurotypical subjects, our QPP based results taken with the broader evidence support a theory of DMN-FPN interaction that points to generally increased DMN-FPN connectivity during rest and internal attention, and decreased DMN-FPN connectivity during externally directed or visual tasks.

#### Ventral attention network

Throughout fMRI literature, evidence characterizes the VAN (also referred to as the salience network or reorienting network) as critical for redirecting attention to salient incoming stimuli (9, 23, 24, 88-90). Across our QPP task spectrum, the VAN exhibited the opposite trend of the DAN and FPN, becoming strongly correlated with the DMN during visual tasks, and more negatively correlated during internally focused tasks. Both categories of visual task in this study (n-back visual working memory and left-right moving dot task) require a high degree of focused external attention and the suppression of distracting stimuli. The VAN becomes more suppressed during external goal-directed tasks, but transiently active when a distracting or novel stimuli is introduced in order to redirect attention (88, 89, 91). Task-negative-like QPP activity for the VAN makes sense then, given that unwanted or poorly tuned reorienting of attention will negatively affect task performance. Increased DMN-VAN alignment during visually demanding tasks matches existing literature from seed-based fMRI analysis. Recent QPP based network analysis from Seeburger et al further corroborates our results, reporting the DMN and VAN to be more positively correlated when task performance was high and more negatively correlated when performance was low (59).

Further exploring aberrant DMN-VAN dynamics could provide a valuable biomarker for attentional disorders (54, 92). Considering seed-based findings of increased DMN-VAN FC in ADHD subjects (92), the aberrant DMN-TPN dynamics reported in Abbas et al (54), and the QPP results in the present study of increased DMN-VAN connectivity during external tasks, it could be the case that ADHD subjects exhibit task-like infraslow brain dynamics (QPPs) even at rest.

### 4.2 Changes to sensory networks (SOM/VIS) across the task-rest spectrum

Sensory networks showed clear shifts across the spectrum of external-internal attention in the group and individual QPP results, indicating that cortical sensory networks participate in dynamic shifting of network relationships on the infraslow QPP timescale. Somatomotor and visual networks showed opposite trends across the spectrum, with SOM becoming much more correlated with the DMN during task, and the VIS becoming more anticorrelated with DMN during task. Increased alignment between the visual network and DAN during a visually demanding task fits an understanding of the DAN as directing top-down goal directed visual attention (10, 21, 27, 93). Likewise, perceptual decoupling is thought to be essential to the maintenance of internal trains of thought, when the DMN is generally most active (44, 77, 94). The need for perceptual decoupling, the separation of intruding somatosensory perception from internal thought processes, might explain why we see high DMN-SOM anticorrelation in the resting state QPPs.

Our results support an updated understanding of sensory areas as being key to cognition and consciousness (95, 96) and not merely evolutionarily ancient substrates of primary sensation (97, 98). Of course, while expansion of the human cortex is disproportionately represented by association areas (2) this does not mean that sensory cortices simply remained static over the course of millions of years of evolution. Rather, sensory networks coevolved with association cortex and underwent selection suited to human behavior (for another good discussion on this topic see “Your Brain Is Not an Onion With a Tiny Reptile Inside”) (99). Dynamic inter-network relationships between the DMN and sensory networks could provide novel insight into our understanding of cortical function as well as a potential biomarker for conditions with altered sensory network activity like autism spectrum disorder (100-102)

In summary, the SOM and VIS networks show strong task related shifts in the QPP, especially in visual tasks, suggesting that dynamic analysis of sensory-motor networks should be included with attentional networks when employing dynamic FC methods to establish future biomarkers.

### 4.4 Limitations

A primary limitation is that the data analyzed in the current study was obtained from a range of different subjects, from different scanners, with slightly different parameters. However, as mentioned in the results section, subjects scanned on the same scanners with identical parameters (HCP data) recapitulated the same task vs rest inter-network changes seen in the other datasets, indicating that the signal of inter-network dynamic changes was greater than noise from dataset-to-dataset variability. Age and sex are factors that have long been known to affect baseline FC (29, 103). While effort was made to include datasets with similar age and sex ratios, these are important biological variables that surely contribute noise to data analyzed here. Demographic variables such as socioeconomic status, a variable that was not included in these datasets, can also have a major impact on FC (38). The inter-network correlation values reported here are on functional time series processed with global signal regression (GSR). There is no consensus on the use of GSR in fMRI preprocessing, with some arguing that GSR increases the possibility of detecting spurious anticorrelations (84, 104). There is also evidence that GSR reveals information in FC patterns missed without GSR (104).

### 4.5 Conclusions

We applied QPP analysis to compare dynamic inter-network correlations between major attentional and sensory networks with the default mode network, a cortical network crucial for neurotypical cognition whose aberrant activity has been implicated in numerous disorders. Using a range of datasets comprising a spectrum of task and rest and external-to-internal attention, we found that DAN, FPN, and VIS networks exhibit task-positive-like responses to visually demanding tasks during QPPs, while the VAN and SOM networks exhibit task-negative-like responses, becoming more aligned with the DMN during visual tasks. These findings based on spatiotemporal dynamics offer additional evidence that the brain does not possess a single task positive or task negative network (105). Rather, cortical networks are in a constant state of dynamic interaction that can flexibly change. The brain is never truly at rest and is constantly integrating external and internal models of reality, a process mediated by the DMN (46, 106). Our results suggest that as the locus of attention shifts from external to internal tasks, cortical network dynamics shift with respect to the DMN in a way that corresponds to the balance of task demands. This is consistent with the literature at large and makes evolutionary sense. The brain is a flexible behavioral system evolved to produce adaptive behavior in the face of constantly shifting environmental variables. It is not surprising then that cortical networks underlying human behavior should themselves show dynamic shifts tracking task demands.

### 4.6 Future Directions

The results presented here provide a reference for comparing altered and pathological cortical dynamics to healthy subjects that can be used to gain insight into disease etiology and to identify diagnostic biomarkers. Specifically, given the evidence of altered DMN-DAN anticorrelation in attentional disorders and mild cognitive impairment, this type of analysis could be applied to datasets of ADHD and Alzheimer’s Disease. The evidence presented here for dynamic shifting of sensory networks with respect to the DMN also suggests that application of this type of QPP analysis to autism spectrum disorder (ASD) could be fruitful, given the altered sensory network activity in subjects with ASD. Considering the growing evidence that the DMN is central to capacities amplified in humans such as theory of mind, advanced language, and the construction of shared neural representations of the world (46), the application of QPP analysis to experimental datasets of subjects engaged in storytelling, problem solving, or economic game theory is also an active area of interest.

## Materials and Methods

All code is Open-Sourced and available online. QPP code referenced in the methods section is available on github: https://github.com/BnzYsf/QPP_Scripts_v0620

Preprocessing code and pipeline are open source and available here: https://fcp-indi.github.io/

All code used for quasi-periodic pattern detection was run in Matlab (Mathworks Inc. 2023) with some plots generated using R studio (Posit team, 2024).

### 2.1 Study design and participants

A range of functional MRI task types were obtained, totaling 327 participants across 788 scans (**Figure 1A**). For a full summary of datasets used, scan type, scans per subject, sources, and the available scan parameters for T1 and functional scans see **Supplemental table 1**. In brief the following task types were used: a left-right moving dot task (107) originally used to measure reaction times in a study of gamers vs. non-gamers; two n-back visual working memory tasks - one from OpenNeuro (108), and one from the Human Connectome Project (109); three types of congruence tasks – Stroop (110), Flanker (111), and Simon (112); a dataset of meditators during resting state and a mindfulness meditation task (83), and a dataset of subjects watching an episode of the Twilight Zone (113).

### 2.3 Quasi-Periodic Pattern Acquisition

Recurring infraslow spatiotemporal patterns (quasi-periodic patterns) were detected using an updated version of the QPP detection algorithm originally employed in rats (50, 114) and more recently in humans (53, 55, 65). For full details on the pattern detection algorithm see the preceding studies. In the present study, a robust version of the pattern algorithm was used that starts at the beginning segment and then iterates through the entire time series, correlating the initial segment with all other segments (**Figure 1C**). Segments which exceed a threshold of correlation (0.2) with the initial QPP template are then averaged. The averaged-updated template is then fed back into the algorithm loop and sliding correlation is repeated until change is negligible between iterations, producing a convergent QPP pattern (65).

Given the established QPP length of approximately 20 seconds in humans (53-56, 59, 60, 65, 115) we selected window lengths for each dataset that resulted in a final QPP length as close to 20 seconds as possible, while still attempting to make adjustments for window lengths that best captured a whole phase of the QPP with minimal inter-QPP phase. The chosen window lengths and resulting final QPP lengths are shown in **Table 1**.

QPPs were detected on a group and individual level. That is, the algorithm was employed on concatenated time series for each group and individually for all 788 scans using the same TR-window length combination from each subject’s respective group. This allowed us to compare inter-network QPP dynamics at both the group and individual level (see **Figure 3**).

### 2.4 Grouping ROIs into networks for inter-network QPP analysis

The output of the QPP detection algorithm is a ∼20 second pattern dominated by reliable and strong anticorrelation between the DMN and TPN. However, as mentioned, the TPN itself is composed of multiple cortical networks depending on the study (DAN, FPN, and/or VAN) (24, 59). And other regions and networks, whose dynamics across task type have not been thoroughly explored, also participate in this strong pattern of anticorrelation (55, 65). To analyze broader network relationships across task types in the QPP and compare them to existing network-based FC findings in the literature from seed-based and dynamic methods, we employed a network-based approach grouping all ROIs into canonical association and sensory networks.

QPP outputs were plotted as 246 ROIs assigned to Yeo’s 7 networks (5), which assigns cortical ROIs into 7 major association and sensory cortices, plus subcortical regions (See supplemental table 2 for a key used to group Brainnetome ROIs into Yeo’s 7 networks). As we were focused primarily on generating an spectrum of inter-network QPP dynamics for the major attentional and sensory networks we excluded the limbic or subcortical networks. The following networks were used: default mode network (DMN), frontoparietal network (FPN), dorsal attention network (DAN), ventral attention network (VAN), somatomotor network (SOM), visual network (VIS) (see Figure 1B for anatomical location of these networks). The task positive network (TPN) is also plotted and included in our analyses but note that this network is just the combination of the DAN and FPN, which show divergent results in our data depending on the task (see figures 3 and 4). As indicated in the following sections, the primary factor driving DMN/TPN anticorrelation across brain states appears to be DMN/DAN anticorrelation, while the DMN/FPN have more flexible correlation.

### 2.5 Measures of inter-network QPP relationships

Waveform plots were generated showing the normalized BOLD signal during the QPP for all networks (Figure 2). Waveform plots were chosen as they allow qualitative inspection of major shifts in network behavior based on task (Figure 2A-F upper panels). Corresponding network correlation plots were generated to capture these shifts quantitatively for each dataset (Figure 2A-F lower panels). Tasks were categorized based on the type of activity. These task categories were reinforced qualitatively based on clear trends we saw in our preliminary results that seemed to indicate a spectrum of inter-network shifts related to external vs internal attention (see Figure 3). This resulted in datasets being categorized as either “visual task” (for visual working memory and the visual reaction time moving dot task), “congruence task” (for congruence tasks such as Flanker, Simon, and Stroop), “rest” (for resting state scans), “meditators rest” (practitioners of mindfulness meditation at rest), “meditation” (same practitioners but during a meditation task), or “Twilight Zone”. Note that Twilight Zone and Meditators are the only datasets in their categories as they did not fit with other tasks categorically or empirically (in terms of their inter-network dynamics, see results).

To generate a spectrum of inter-network DMN dynamic fingerprints as a potential biomarker, focused analysis was made with respect to the DMN (Figures 4-5). Initially, the correlation between all networks and the DMN was calculated at the group level (Figure 3A). As notable differences immediately became apparent across the task spectrum in several networks, we repeated all DMN-to-network correlations at the individual level for comparison (Figure 3 A-B). DMN-to-network correlation from individual QPPs (Figure 3B) was used for statistical comparison (Figure 3C), as group level correlations lack variance and mean. Scans were grouped by task category to test differences between “visual task”, “congruence task” and “rest” QPPs. Note that meditator and Twilight Zone groups were excluded from statistical comparison as they have much smaller sample sizes and showed empirically different trends from the other categories. These miscellaneous groups show interesting trends however, informative for interpreting task vs rest and for guiding future experiments, and were therefore included for visual comparison. Statistical comparisons of DMN-to-network correlations (Figure 3C) were made using the Kruskal-Wallis test for multiple comparisons. All correlation values were Fisher transformed before statistical comparison to account for non-normal distribution. Multiple comparison correction was done using the conservative Bonferroni method (116) and assuming 7 comparisons between the 8 canonical networks used. (Standard α = .05 was adjusted for 7 comparisons (.05/7) to α = .007). To capture the idea of DMN-to-network correlations in the QPP as a “fingerprint” that can be used as a neurotypical reference to compare to future studies, we generated a DMN-to-network QPP fingerprint for each category of scan (Figure 4). The qualitative differences between infraslow QPP fingerprints are discussed in the results section, while the underlying statistical differences are captured network by network in Figure 3C.

## Acknowledgments

We would like to acknowledge the Dhamala lab at Georgia State University and Dr. Jordan for collaborating and making available their moving dot task data. We also thank Dr. Wendy Hasenkamp and Dr. Larry Barsalou for sharing their meditation scans with us. Finally, we want to acknowledge that this study was made possible by OpenNeuro, the Human Connectome Project, and CABI, for promoting large scale open data sharing of fMRI scans. This research was funded by grants from the National Institutes of Health: 1R01Ns078095, 1R01AG062581, 1R01EB029857.

---

**Supplementary Table 1.** Scan sources and parameters.

| Dataset | Source: | T1 Anatomical | Functional Scan |
| --- | --- | --- | --- |
| **Video Gamer Task (VG)**<br>N=47, 4 task scans per subject | Dr. Mukeshwar Dhamala (Georgia State University) and Timothy Jordan (UCLA). Georgia State/Georgia Tech Center for Advanced Brain Imaging (CABI). | 3T S Magnetom Prisma MRI scanner; TR=2530ms; TE1-4: 1.69–7.27ms; flip angle:7°;Voxel Size:1mm<sup>3</sup> | 3T S Magnetom Prisma MRI scanner; TR=535ms; TE=30ms; flip angle:46°; FOV:240mm; Voxel Size: 3.8x3.8x4mm; slices:32 |
| **Visual Working (VWM)**<br>Memory Task<br>N=19 (13M 6F)<br>Mean Age: 22.4, 3 task scans per subject | OpenNeuro<br>(Hampshire, Adam and Soreq, Eyal (2019) | Siemens 3T MRI TrioTim scanner; TR=2500ms; TE=2.9ms; flip angle: 9°; Voxel Size:1mm<sup>3</sup> | Siemens 3T MRI TrioTim scanner; TR=2000ms; TE=30ms; flip angle:45°; FOV:240mm; Voxel Size: 3mm<sup>3</sup>; slices:36 |
| **Simon Task**<br>N=21, 2 task scans per subject | OpenNeuro<br>(Kelly AMC and Milham MP (2018). Simon task. OpenNeuro). | Siemens Allegra 3T scanner; TR=2500ms; TE=3.93ms; flip angle:8°; FOV:256mm; Voxel Size:1mm<sup>3</sup> | Siemens Allegra 3T scanner; TR=2000ms; TE=30ms; flip angle:80°; FOV:192mm; Voxel Size: 3x3x4mm; slices:40 |
| **Stroop Task**<br>N=28 (20M 10F)<br>Mean Age: 31yr, 1 task scan per subject | OpenNeuro<br>(Timothy D. Verstynen (2018). Stroop Task. OpenNeuro). | Siemens Verio 3T scanner | Siemens Verio 3T scanner; TR=1500ms; TE=20ms; flip angle:90°; FOV:240mm; Voxel Size: 3.5mm isotropic; slices:30 |
| **Flanker Task**<br>N=26, 2 task scans per subject | OpenNeuro<br>(Kelly AMC and Uddin LQ and Biswal BB and Castellanos FX and Milham MP (2018). Flanker task (event-related). OpenNeuro. | Siemens Allegra 3T scanner; TR=2500ms; TE=3.93ms; flip angle:8°; FOV:256mm; Voxel Size:1mm<sup>3</sup> | Siemens Allegra 3T scanner; TR=2000ms; TE=30ms; flip angle:80°; FOV:192mm; Voxel Size: 3x3x4mm; slices:40 |
| **CABI Rest**<br>N=99<br>Mean Age: 26yr, 1 rest scan per subject, 2 rest scans for most subjects | Dr. Eric Schumacher, Georgia Institute of Technology. Georgia State/Georgia Tech Center for Advanced Brain Imaging (CABI). | Siemens 3T Trio MRI scanner; TR=2250ms; TE=3.98ms; flip angle:9°; FOV:256mm; Voxel Size:1mm<sup>3</sup> | Siemens 3T Trio MRI scanner; TR=2000ms; TE=30ms; flip angle:90°; FOV:240mm; Voxel Size: 3x3x3mm; slices:37 |
| **HCP Rest/Task**<br>N = 50<br>Ages 26-36, 2 rest and 2 | Human Connectome Project. Washington University. (Van Essen et al., 2012). | Siemens Skyra 3T; 3D MPRAGE sequence; TR = 2400 ms, TE = 2.14 ms, TI = 1000 ms, FA = 8°, | Siemens Skyra 3T; Gradient-echo Echo Planar Imaging; TR = 720 ms, TE = 33.1 ms, FA = 52°, FOV = 208 mm × 180 mm |
| task scans<br>per subject |  | FOV = 224 mm × 224 mm,<br>voxel size 0.7 mm<sup>3</sup> | (RO x PE), matrix = 104 × 90<br>(RO x PE), slice<br>thickness = 2.0 mm; 72<br>slices; 2.0 mm isotropic<br>voxels, multi-band factor = 8,<br>echo spacing 0.58 ms |
| **Meditators**<br>N=14 (3M<br>11F)<br>Ages 28-66,<br>1 rest and 1<br>meditation<br>task scan<br>per subject | Dr. Wendy Hasenkamp and Dr. Larry<br>Barsalou, Dr. Christine Wilson-Mendenhall,<br>Emory University. (Hasenkamp et al.,<br>2012). | Siemens 3T Trio MRI<br>scanner; TR=2600ms;<br>TE=3.9ms; FOV:240mm | Siemens 3T Trio MRI<br>scanner; TR=1500ms;<br>TE=30ms;<br>flip angle:90°; FOV:192mm;<br>Voxel Size: 4x3x3mm;<br>slices:18 |
| **Twilight**<br>**Zone**<br>N=24, 1 task<br>(watching<br>Twilight<br>Zone) per<br>subject | OpenNeuro. (J. Chen and C. J. Honey and<br>E. Simony and M. J. Arcaro and K. A.<br>Norman and Hasson, U. (2019). Twilight<br>Zone Movie Watching Dataset. OpenNeuro.<br>doi:<br>10.18112/openneuro.ds001145.v1.0.0). | Siemens 3T Skyra scanner;<br>Voxel Size:0.89mm<sup>3</sup> | Siemens 3T Skyra scanner;<br>TR=1500ms; TE=28ms;<br>flip angle:64°; FOV:192mm;<br>Voxel Size: 4x3x3mm;<br>slices:27 |

**Supplementary Table 2.** Brainnetome coordinates of ROIs assigned to each network.

| DMN | DAN | FPN | VAN | VIS | SOM |
| --- | --- | --- | --- | --- | --- |
| 3 | 7 | 1 | 2 | 105 | 9 |
| 5 | 8 | 4 | 15 | 106 | 10 |
| 6 | 25 | 12 | 37 | 108 | 53 |
| 11 | 26 | 16 | 38 | 112 | 54 |
| 13 | 30 | 17 | 39 | 113 | 57 |
| 14 | 55 | 18 | 40 | 114 | 58 |
| 23 | 56 | 19 | 61 | 119 | 59 |
| 33 | 63 | 20 | 62 | 120 | 60 |
| 34 | 64 | 21 | 65 | 135 | 66 |
| 35 | 85 | 22 | 123 | 136 | 67 |
| 41 | 86 | 24 | 124 | 151 | 68 |
| 42 | 91 | 28 | 167 | 152 | 71 |
| 43 | 92 | 29 | 168 | 182 | 72 |
| 44 | 97 | 31 | 169 | 189 | 73 |
| 51 | 98 | 32 | 170 | 190 | 74 |
| 52 | 107 | 36 | 173 | 191 | 75 |
| 79 | 125 | 46 | 174 | 192 | 76 |
| 80 | 126 | 82 | 180 | 193 | 131 |
| 81 | 127 | 99 | 183 | 194 | 132 |
| 83 | 128 | 100 | 184 | 195 | 145 |
| 84 | 129 | 137 | 185 | 196 | 146 |
| 87 | 130 | 138 | 186 | 197 | 149 |
| 88 | 133 | 142 |  | 198 | 155 |
| 95 | 134 | 147 |  | 199 | 156 |
| 121 | 139 | 148 |  | 200 | 157 |
| 122 | 140 | 166 |  | 202 | 158 |
| 141 | 143 |  |  | 203 | 160 |
| 144 | 150 |  |  | 204 | 161 |
| **153** | 159 |  |  | 205 | 162 |
| 154 | 201 |  |  | 206 | 163 |
| **175** |  |  |  | 207 | 164 |
| 176 |  |  |  | 208 | 171 |
| 179 |  |  |  | 209 | 172 |
| 181 |  |  |  | 210 |  |
| 187 |  |  |  |  |  |
| 188 |  |  |  |  |  |

